# A single dose of ChAdOx1 MERS provides broad protective immunity against a variety of MERS-CoV strains

**DOI:** 10.1101/2020.04.13.036293

**Authors:** Neeltje van Doremalen, Elaine Haddock, Friederike Feldmann, Kimberly Meade-White, Trenton Bushmaker, Robert J. Fischer, Atsushi Okumura, Patrick W. Hanley, Greg Saturday, Nick J. Edwards, Madeleine H.A. Clark, Teresa Lambe, Sarah C. Gilbert, Vincent J. Munster

## Abstract

Middle East respiratory syndrome coronavirus (MERS-CoV) continues to infect humans via the dromedary camel reservoir and can transmit between humans, most commonly via nosocomial transmission. Currently, no licensed vaccine is available. Previously we showed that vaccination of transgenic mice with ChAdOx1 MERS, encoding the MERS S protein, prevented disease upon lethal challenge. In the current study we show that rhesus macaques seroconverted rapidly after a single intramuscular vaccination with ChAdOx1 MERS. Upon MERS-CoV challenge vaccinated animals were protected against respiratory injury and pneumonia and had a reduction in viral load in lung tissue of several logs. Furthermore, we did not detect MERS-CoV replication in type I and II pneumocytes of ChAdOx1 MERS vaccinated animals. A prime-boost regimen of ChAdOx1 MERS boosted antibody titers, and viral replication was completely absent from the respiratory tract tissue of these rhesus macaques. Finally, we investigated the ability of ChAdOx1 MERS to protect against six different MERS-CoV strains, isolated between 2012 to 2018, from dromedary camels and humans in the Middle East and Africa. Antibodies elicited by ChAdOx1 MERS in rhesus macaques were able to neutralize all MERS-CoV strains. Vaccination of transgenic hDPP4 mice with ChAdOx1 MERS completely protected the animals against disease and lethality for all different MERS-CoV strains. The data support further clinical development of ChAdOx1 MERS supported by CEPI.

**One Sentence Summary:** Prime-only vaccination with ChAdOx1 MERS provides protective immunity against HCoV-EMC/2012 replication in rhesus macaques, and a wide variety of MERS-CoV strains in mice.

## Introduction

Middle East respiratory syndrome coronavirus (MERS-CoV) was first identified in 2012 and has since infected >2400 people. The clinical spectrum of MERS-CoV infection in humans varies from asymptomatic to severe respiratory disease and death. Patients present with influenza-like symptoms such as a fever and shortness of breath. Thereafter, they may develop pneumonia and may require mechanical ventilation and support in an intensive care unit. The case-fatality rate of MERS-CoV infection is currently 34.5% (*1*).

Human-to-human transmission of MERS-CoV is relatively limited and occurs mainly in nosocomial settings but has been reported in local communities as well (*2*). In 2015, a traveler from the Middle East to South Korea caused an outbreak involving 186 people and 38 fatalities (*3*). The nosocomial outbreak lasted from May to July, and 16,752 people were isolated with MERS-CoV-like symptoms. At least three superspreaders were identified in the outbreak, who infected 28, 85 and 23 patients respectively (*4*). The introduction and spread of MERS-CoV in South Korea underscores the potential of this virus to cause epidemics outside of the Arabian Peninsula. MERS-CoV has continued to cause disease in humans and so far in 2019, 199 cases have been reported in the Kingdom of Saudi Arabia (KSA), of which 24% were fatal (*1*). The ongoing circulation of MERS-CoV and subsequent outbreaks in the human population highlight the need of an efficient MERS-CoV vaccine.

MERS-CoV is mainly prevalent in the Arabian Peninsula, with the majority of cases occurring in KSA (84%) (*5*). However, phylogenetically diverse MERS-CoV strains have been isolated from Africa and the Middle East (*6-8*) and antigenic differences have been reported between spike (S) proteins, the main antigen utilized in MERS-CoV vaccines, from the Middle East and Africa (*7*). Thus, it is important that a MERS-CoV vaccine is protective against a variety of diverse MERS-CoV strains.

Thus far, only four vaccines have been tested in rhesus macaques. These are a DNA vaccine (*9*), a vaccine based on the receptor binding domain (RBD) of the MERS-CoV S adjuvanted with alum (*10*), a combination of S plasmid DNA and S1 protein (*11*), and virus-like particles based on MERS-CoV S, matrix, and envelope proteins (*12*). Of the four vaccine studies, only three studies included challenge studies, all of which used MERS-CoV isolates from 2012. To date there have been no reports of efficacy of a single-dose MERS-CoV vaccine in NHPs.

We recently demonstrated that vaccination of mice with a replication-deficient simian adenovirus vaccine vector (ChAd) encoding full-length MERS-CoV S protein (ChAdOx1 MERS) elicited high-titer MERS-CoV neutralizing antibodies and a robust CD8+ T cell response against the S protein (*13*). In addition, ChAdOx1 MERS vaccination resulted in full protection of human DPP4 transgenic mice against a lethal challenge with MERS-CoV (*14*). ChAdOx1 MERS vaccination of dromedary camels was immunogenic and reduced MERS-CoV shedding after challenge in a natural transmission model (*15*). ChAd-vectored vaccines against malaria, HIV, influenza, hepatitis C, tuberculosis, Ebola, and others show an excellent immunogenicity and safety profile in humans. In the current manuscript, we show that a single dose of ChAdOx1 MERS vaccine protects rhesus macaque model against a mucosal challenge with HCoV-EMC/2012. Serum obtained from vaccinated rhesus macaques was able to neutralize six diverse MERS-CoV strains. Furthermore, a single dose of ChAdOx1 MERS vaccine protects hDPP4 transgenic mice against all evaluated MERS-CoV strains.

## Results

### ChAdOx1 MERS elicits a productive immune response in rhesus macaques

Three groups of six animals each were vaccinated with ChAdOx1 MERS via a prime-boost regimen (−56 DPI and -28 DPI) or prime-only regimen (−28 DPI) or with ChAdOx1 GFP (−56 DPI and -28 DPI). Animals were then challenged with 7 x 10^6^ TCID_50_ HCoV-EMC/2012 on 0 DPI via combined intratracheal, intranasal, oral and ocular route (*16*). Blood samples were taken at days of vaccination, 14 days post each vaccination, and at challenge (*Figure 1A*). The humoral immune response to vaccination was examined by ELISA using S protein as well as by virus neutralization assay. S protein-specific IgG antibodies were detected as early as 14 days post vaccination in both ChAdOx1 MERS vaccinated groups. All animals in these groups had S-protein-specific antibodies at 0 DPI, whereas no antibodies against S protein were found before vaccination, or at any time in the ChAdOx1 GFP vaccinated group (*Figure 1B*). Neutralizing antibodies were detected in 11 out of 12 ChAdOx1 MERS vaccinated animals at time of challenge. One animal in the prime-only vaccination group did not have detectable neutralizing antibodies at this time and had the lowest antibody titer as measured by ELISA (1600) (*Figure 1C*). A second ChAdOx1 MERS vaccination at -28 DPI resulted in a statistically significant increase in S protein-specific ELISA titer (mean titer -28 DPI = 1867; 0 DPI = 10,667; P<0.0001) and neutralizing antibody titer (mean titer -28 DPI = 28; 0 DPI = 160; P<0.0001) as determined via Tukey’s multiple comparisons test, even though neutralizing antibodies against the ChAdOx1 vector could be detected at the time of the second vaccination (*Figure S1A*). This suggests that the presence of neutralizing antibodies against ChAdOx1 does not prevent the vaccine vector from boosting the immune response.

**Fig. 1.**
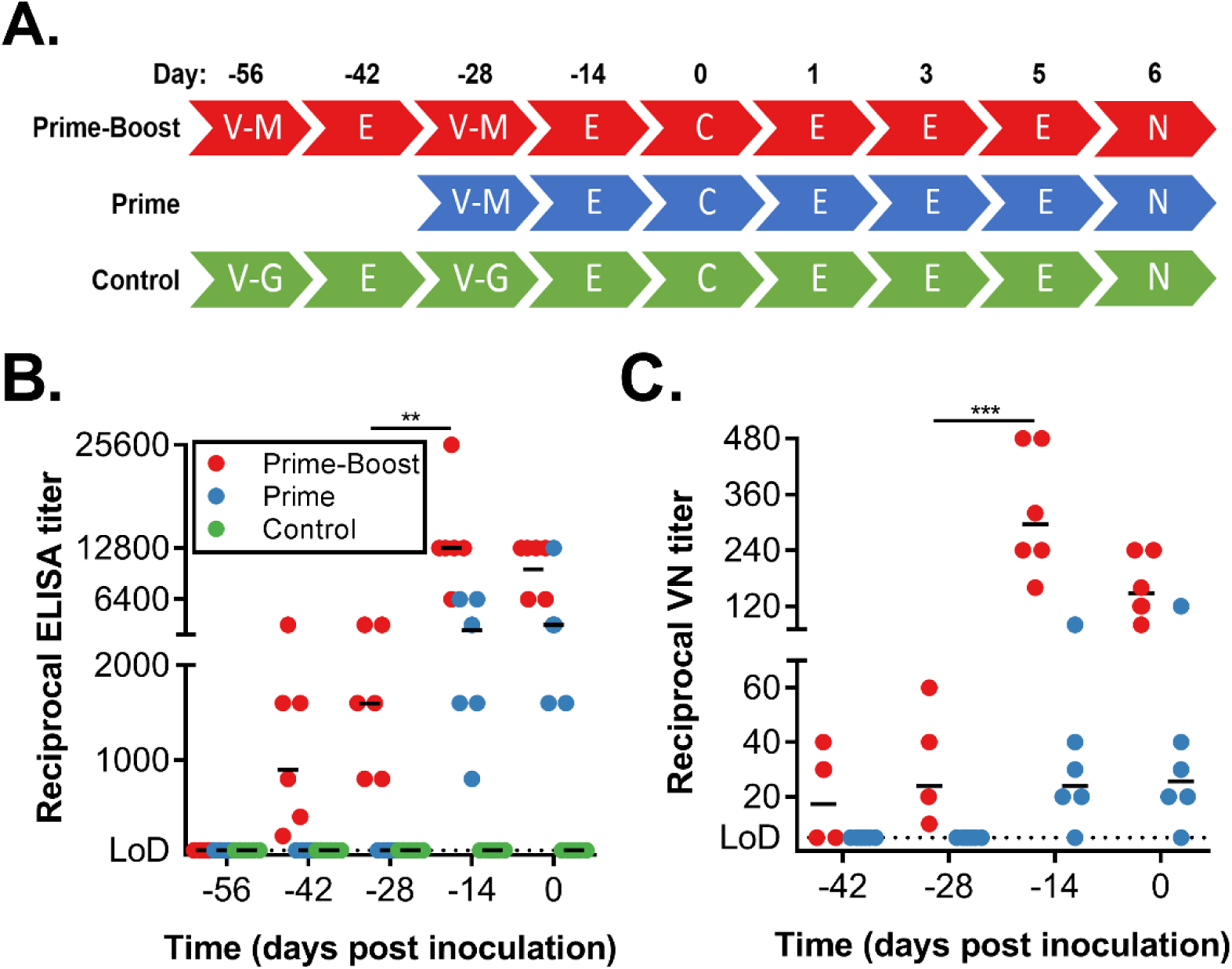
Vaccination of rhesus macaques with ChAdOx1 MERS elicits a humoral immune response. Serum samples were collected from non-human primates at times of vaccination (−56 DPI and -28 DPI), 14 days later and at challenge. **(A)** Overview of experimental timeline. V-M = vaccination with ChAdOx1 MERS; V-G = vaccination with ChAdOx1 GFP; E = exam; C = challenge and exam; N = exam and necropsy **(B)** Two-fold serial-diluted serum samples were tested for MERS-CoV S-specific antibodies using ELISA. **(C)** ELISA-positive two-fold serial-diluted serum samples were tested for neutralizing antibodies against MERS-CoV in VeroE6 cells. Line = geometric mean, dotted line = limit of detection. Statistical significance between -28 DPI and -14 DPI in the prime-boost group was determined via one-tailed paired Student’s t-test. ** = p-value<0.01; *** = p-value<0.001

### Vaccination with ChAdOx1 MERS reduces disease severity

Animals vaccinated with ChAdOx1 GFP and challenged with MERS-CoV showed similar clinical signs as previously reported (*16*), such as a decreased appetite and increased respiratory rate. Clinical signs were scored using a standard non-human primate scoring sheet, focusing on areas such as general appearance, nasal discharge and food intake. On average, ChAdOx1 MERS vaccinated animals had a lower clinical score than ChAdOx1 GFP vaccinated animals (*Figure 2A*). All animals underwent exams on 0, 1, 3, 5, and 6 DPI. No significant changes in weight or body temperature were observed for the duration of the study. Peripheral capillary oxygen saturation (spO_2_) measurements supported a decrease in oxygen saturation from the baseline in ChAdOx1 GFP vaccinated animals, but not in ChAdOx1 MERS vaccinated animals (*Figure 2B*).

**Fig. 2.**
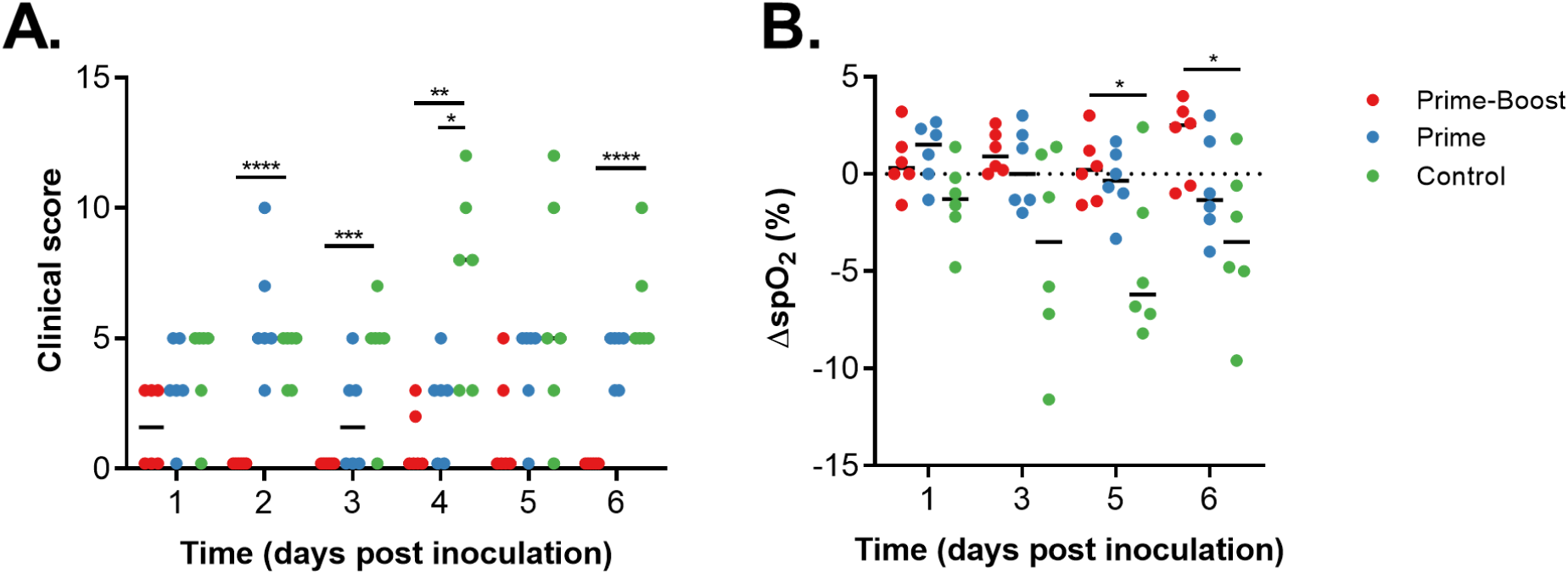
Clinical scoring and spO_2_ values are improved in ChAdOx1 MERS vaccinated animals compared to ChAdOx1 GFP vaccinated animals. **(A)** Animals were evaluated daily and clinical score assessed using an established scoring sheet. **(B)** Changes in oxygen saturation from pre-inoculation values (Δ% spO2) were determined on exam days. Statistical significance between groups was determined via two-tailed unpaired Student’s t-test. Line = median; dotted line = baseline value; * = p-value<0.025; ** = p-value<0.01; *** = p-value<0.001; **** = p-value<0.0001.

Ventrodorsal and lateral thoracic radiographs were collected on all exam days. All animals vaccinated with ChAdOx1 GFP showed moderate to severe pulmonary interstitial infiltrations, whereas animals vaccinated with ChAdOx1 MERS showed only mild signs of respiratory disease. A severe collapsed lung was observed on 3 DPI in two animals in the prime-boost group, likely caused by the bronchoalveolar lavage (BAL) performed on 1 DPI. Full details of all observed signs can be found in Table S2. (*Figure 3A, Table S2*). All lung lobes (right cranial, right middle, right caudal, left cranial, left middle, left caudal) of each individual animal were scored for severity of disease signs for each day radiographs were taken, and average scores were compared. Scores obtained from animals vaccinated with ChAdOx1 GFP were significantly higher from animals vaccinated with a prime-boost regimen of ChAdOx1 MERS on 3, 5, and 6 DPI (*Figure 3B*).

**Fig. 3.**
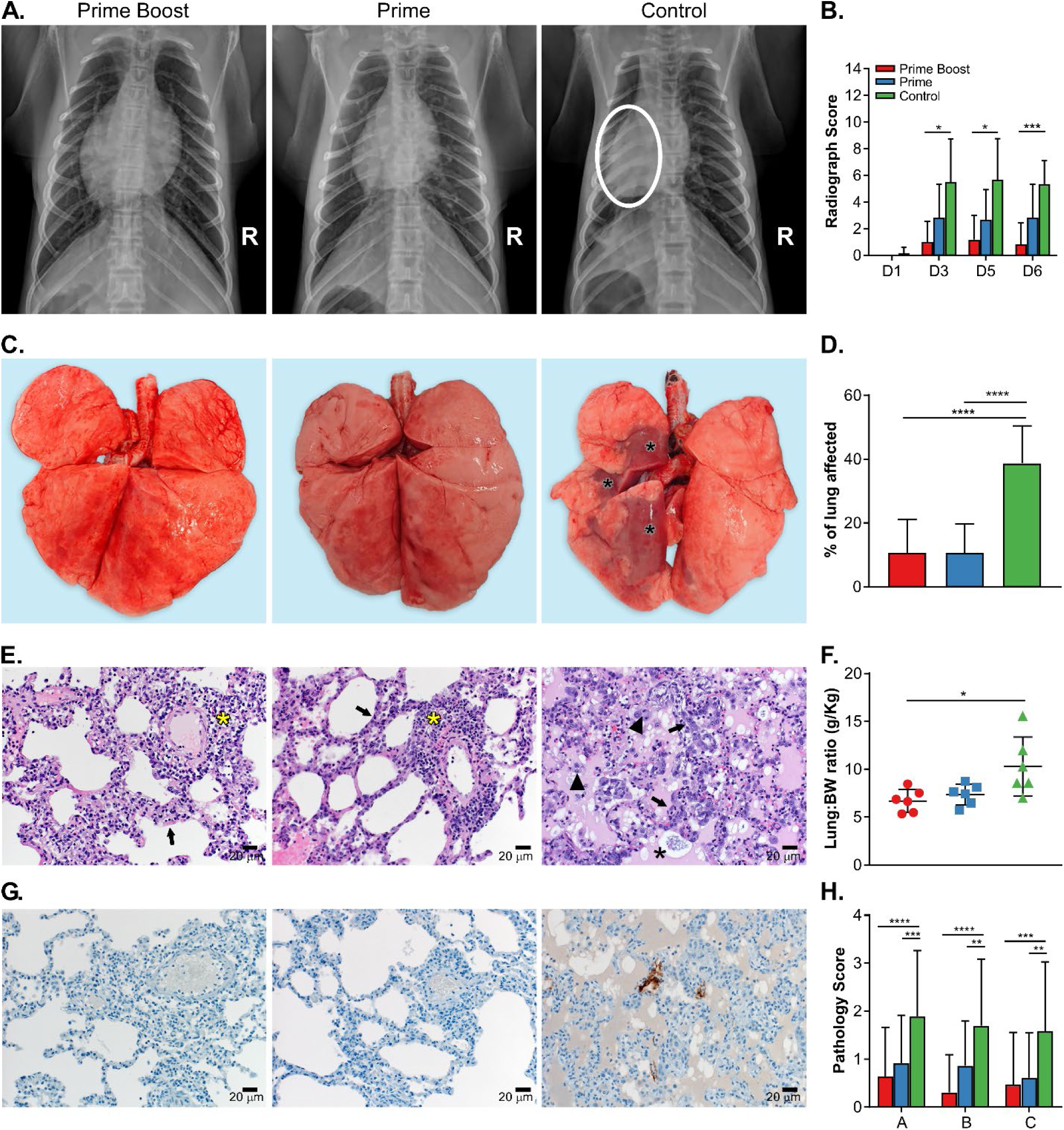
Single-dose vaccination with ChAdOx1 MERS protects rhesus macaques against bronchointerstitial pneumonia caused by MERS-CoV challenge. Rhesus macaques were vaccinated with a prime-boost or prime-only regimen of ChAdOx1 MERS, or with ChAdOx1 GFP and challenged with MERS-CoV. **(A)** Ventrodorsal thoracic radiographs collected on 6 DPI. A marker (R) indicates right side of animal. No pathologic changes were observed in animals vaccinated with ChAdOx1 MERS via a prime-boost or prime-only regimen. Animal vaccinated with ChAdOx1 GFP shows focally extensive area of increased pulmonary opacity and deviation of the cardiac silhouette, highlighted in the circle located in the middle and caudal lung lobes. **(B)** Thoracic radiographs of each animal were scored per lung lobe resulting in a maximum score of 18. Values were averaged per group per day, shown is mean with standard deviation. See Table S2 for more details. **(C)** Gross pathology of lungs shows no pathologic changes in ChAdOx1 MERS vaccinated animals, and focally extensive areas of consolidation in left cranial, middle and caudal lung lobes in control animals (asterisks). **(D)** Gross lung lesions were scored for each lung lobe, ventral and dorsal. Values were averaged per group, shown is mean with standard deviation. **(E)** Lung tissue sections were stained with hematoxylin and eosin. Moderate numbers of lymphocyte accumulation around pulmonary arterioles (asterisks), and mild thickening of alveolar septae by lymphocytes and macrophages (arrow) in lung tissue of animals vaccinated with ChAdOx1 MERS. Marked bronchointerstitial pneumonia with abundant pulmonary edema and fibrin (asterisks), type II pneumocyte hyperplasia (arrow), and increased numbers of alveolar macrophages (arrowhead) in lung tissue of control animals. Magnification = 200x. **(F)** Lung to body weight ratio was determined for all animals at necropsy. Shown is mean with standard deviation. **(G)** Lung tissue sections were stained with antibody against MERS-CoV antigen, which is visible as a red-brown staining. No immunoreactivity was found in ChAdOx1 MERS vaccinated animals, whereas multifocal immunoreactivity of type I and II pneumocytes could be found in lung tissue of ChAdOx1 GFP vaccinated animals. **(H)** Lung tissue sections were scored on severity of lesions (0=no lesions, 1=1-10%, 2=11-25%, 3=26-50%, 4=51-75%, 5=76-100%) and averaged per group. Shown is mean with standard deviation. A = bronchointerstitial pneumonia; B = type II pneumocyte hyperplasia; C = Hemorrhages, edema, fibrin deposits. Statistical significance between groups was determined via two-tailed unpaired Student’s t-test. * = p-value<0.025; ** = p-value<0.01; *** = p-value<0.001; **** = p-value<0.0001.

Upon necropsy on 6 DPI, gross lung lesions were more prevalent in animals vaccinated with ChAdOx1 GFP than animals vaccinated with ChAdOx1 MERS (*Figure 3C-D*). Animals vaccinated with ChAdOx1 GFP showed focally extensive areas of consolidation in all lung lobes and lungs generally failed to collapse. Mediastinal lymph nodes were often edematous and enlarged. Animals that received a vaccination with ChAdOx1 MERS, either via a prime-boost or prime-only regimen, had either no lesions or limited small multifocal areas of consolidation and congestion.

An increased lung:body weight ratio is an indicator of pulmonary edema. Animals in the control group had significantly higher lung:body weight ratios compared to ChAdOx1 MERS vaccinated animals and there was minimal difference between prime-boost and prime-only ChAdOx1 MERS vaccinated animals (*Figure 3F*).

Lung tissue sections were stained with hematoxylin and eosin or with MERS-CoV specific antibodies. All slides were evaluated by a board-certified veterinary pathologist blinded to study group allocations. In animals which received a vaccination with ChAdOx1 MERS, either prime-boost or prime-only, a minimal to mild bronchointerstitial pneumonia was present characterized by mild thickening of alveolar septae by lymphocytes and macrophages. Pulmonary vessels were bound by moderate numbers of lymphocytes. In stark contrast, in lung tissue obtained from animals vaccinated with ChAdOx1 GFP, moderate to marked bronchointerstitial pneumonia was present throughout the lung lobes characterized by thickening of the alveolar septae by lymphocytes and macrophages, edema and fibrin. Alveoli contained abundant edema and fibrin and moderate to abundant numbers of alveolar macrophages, neutrophils and necrotic debris. Inflammation often surrounded bronchioles and pulmonary vasculature and type II pneumocyte hyperplasia was prominent. Additionally, presence of MERS-CoV antigen by immunohistochemistry was found only in lungs of animals vaccinated with ChAdOx1 GFP within type I & II pneumocytes and was not found in lung tissue of ChAdOx1 MERS vaccinated animals. Severity of bronchointerstitial pneumonia, type II pneumocyte hyperplasia and hemorrhages, edema, and fibrin deposits was scored. Statistically significant differences between animals vaccinated with ChAdOx1 MERS or ChAdOx1 GFP were found for all three categories. (*Figure 3E, F, H*). Thus, vaccination with ChAdOx1 MERS, either via a prime-boost regimen or prime-only regimen, significantly decreased the severity of pulmonary pathology and protected rhesus macaques against bronchointerstitial pneumonia.

### Vaccination with ChAdOx1 MERS limits virus replication in the respiratory tract

BAL was performed on all animals on 1, 3, 5 and 6 DPI and the amount of viral RNA, mRNA and infectious virus was determined. In the prime-boost group, infectious virus was only detected at 1 DPI (N=3) after inoculation, and in the prime-only group at 1 DPI (N=6) and 3 DPI (N=3) but not thereafter. In contrast, infectious virus was detected on all days in BAL fluid from the control group (*Figure 4A*). Viral mRNA in BAL fluid of animals in the prime-boost group was only detected at 1 DPI (N=5) and 3 DPI (N=1). In contrast, viral mRNA in BAL fluid from animals who received a single vaccination with ChAdOx1 MERS mRNA was detected on all days, but the amount was reduced compared to animals vaccinated with ChAdOx1 GFP (*Figure S1B*). Viral RNA as measured by UpE qRT-PCR assay could be detected in all groups up to 6 DPI. However, the number of genome copies/mL detected was lower in animals vaccinated with ChAdOx1 MERS compared to animals vaccinated with ChAdOx1 GFP (*Figure 4A*). A significant association was found between higher ELISA titer or VN titer and lower levels of viral RNA, mRNA or infectious virus in BAL fluid for all days, except 6 DPI for infectious virus (Spearman’s rank correlation coefficient, *Table S2*).

**Fig. 4.**
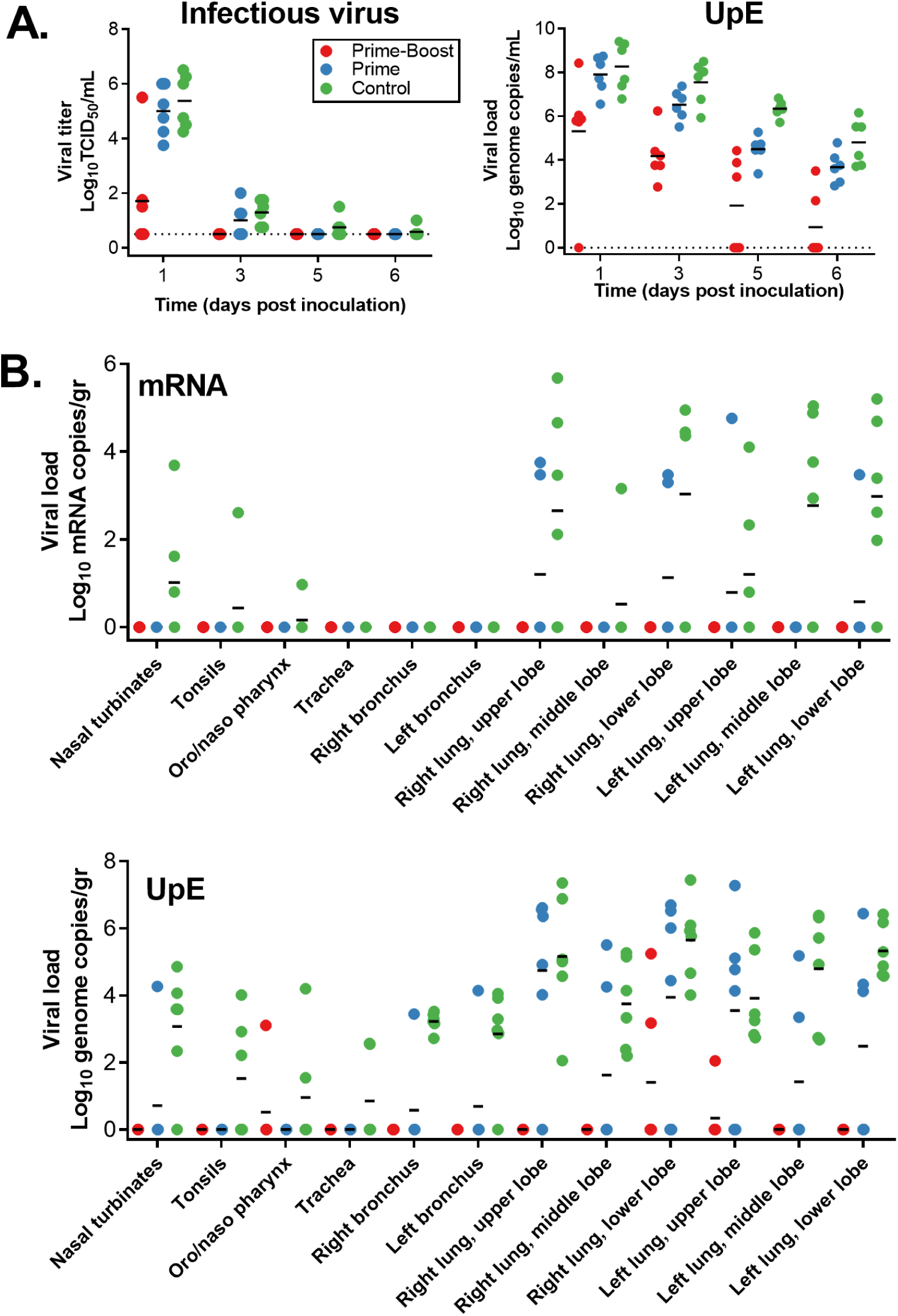
Vaccination with ChAdOx1 MERS results in reduced virus replication in the respiratory tract. **(A)** Infectious virus titers and viral load were determined in BAL fluid. Individual values are depicted. **(B)** UpE and mRNA copies were determined in respiratory tract tissues collected at 6 DPI. Individual values are depicted. Line = geometric mean, dotted line = limit of detection.

All animals were euthanized on 6 DPI and tissues were analyzed for the presence of viral RNA, mRNA or infectious virus. Infectious virus titers were only found in nasal turbinate tissue (N=1) and lung lobe tissue (N=4) from control animals. No viral mRNA was found in all tissues obtained from animals that received a prime-boost regimen. In tissue from animals that received a prime-only regimen, limited viral mRNA could be found in upper and lower lung lobe tissues (N=4). In contrast, mRNA could be found in respiratory tract tissues of all control animals, as well as in conjunctiva (*Figure 4B*). Viral RNA was detected in tissues from all groups but was mainly found in lung lobes and bronchi. Viral load was higher for lower respiratory tract tissue obtained from animals vaccinated with ChAdOx1 GFP (N=6) than from animals receiving a prime-only (N=6) or a prime-boost regimen of ChAdOx1 MERS (N=2), (*Figure 4B*).

### Cytokines are upregulated in lung tissue of ChAdOx1 GFP vaccinated animals compared to ChAdOx1 MERS vaccinated animals

The presence of 23 cytokines was evaluated in lung tissue. Several cytokines were upregulated in animals vaccinated with ChAdOx1 GFP compared to animals vaccinated with ChAdOx1 MERS, including IL2, IL6, IL8, IL18, MCP-1, MIP-1a, and TGF-α (*Figure 5*). This is indicative of a local enhanced immune response in animals vaccinated with ChAdOx1 GFP, but not in animals vaccinated with ChAdOx1 MERS.

**Fig. 5.**
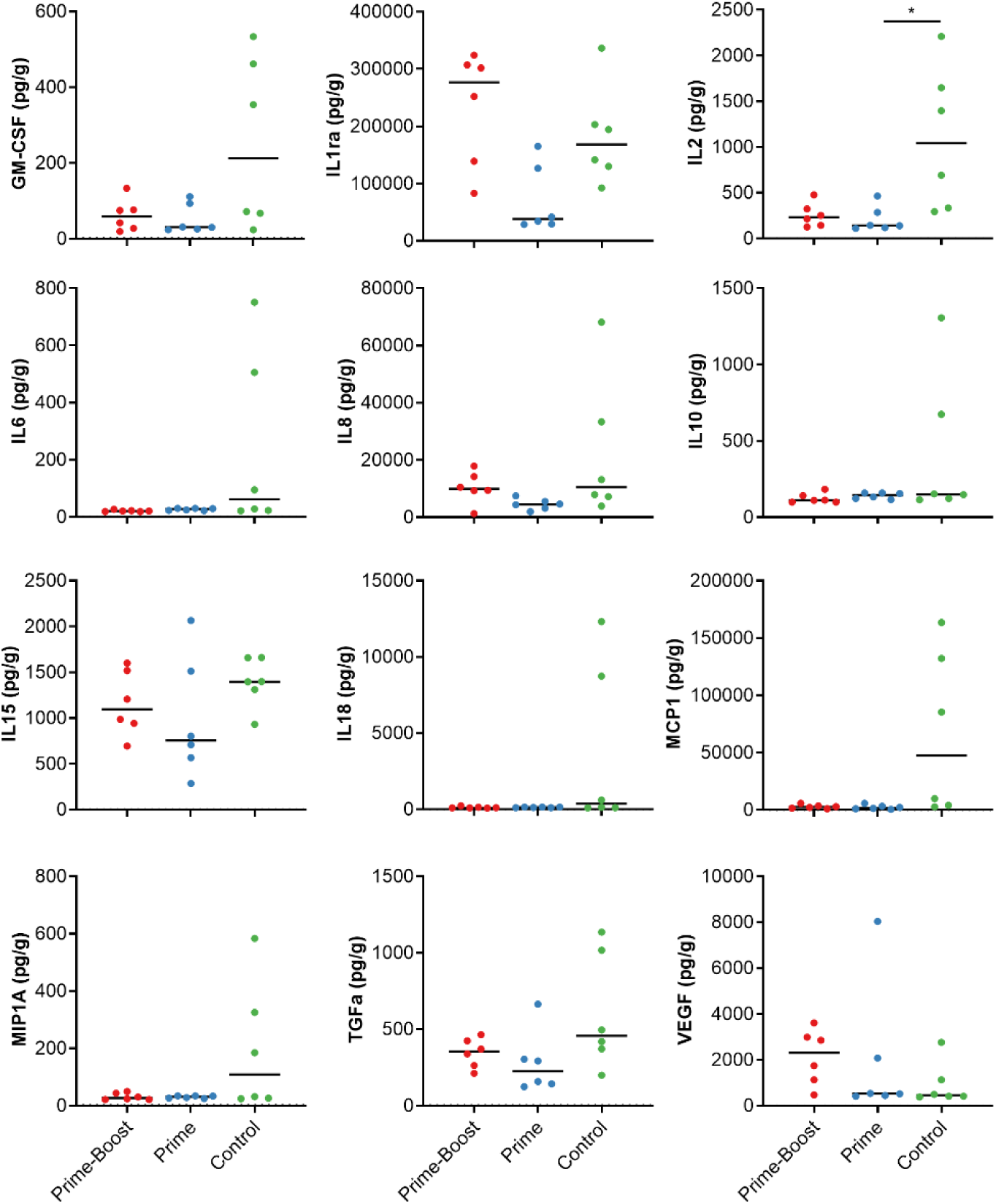
Several cytokines are upregulated in lung tissue of animals vaccinated with ChAdOx1 GFP compared to animals vaccinated with ChAdOx1 MERS. Cytokines in homogenized lung tissue were detected using a 23-plex NHP panel on the Luminex. Cytokines with no values over the LOD were excluded. Line = median; * = p-value<0.025.

### Antibodies elicited by ChAdOx1 MERS vaccination neutralize different MERS-CoV strains

We selected six different strains of MERS-CoV (*Figure S2*) and tested neutralizing capability of serum obtained at 0 DPI from animals vaccinated with a prime-boost regimen. Strains were selected based on geographical location (Saudi Arabia, South Korea, and Burkina Faso), host (dromedary camel or human) and time of isolation (2012 to 2018). All six strains were neutralized by antibodies elicited by ChAdOx1 MERS vaccination (*Table 1*).

**Table 1.** Neutralizing titer of serum obtained from animals vaccinated with a prime-boost regimen of ChAdOx1 MERS against different MERS-CoV strains.

| Full virus name | Abbreviation | Genbank accession number | Animal number |  |  |  |  |  |
| --- | --- | --- | --- | --- | --- | --- | --- | --- |
|  |  |  | 1 | 2 | 3 | 4 | 5 | 6 |
| Hu/HCoV/EMC/2012 | EMC/12 | JX869059 | 120 | 480 | 120 | 240 | 240 | 120 |
| Hu/Saudi Arabia/Rs924/2015 | KSA/15 | KY688119 | 240 | 160 | 80 | 160 | 120 | 60 |
| Hu/Korea/Seoul/SNU1-035/2015 | SK/15 | KU308549 | 160 | 120 | 60 | 160 | 60 | 60 |
| Riyadh/KSA-18013832/2018 | KSA/18 | MN723544 | 320 | 320 | 160 | 480 | 120 | 80 |
| Camel/Saudi Arabia/KFU-HKU1/2013 | C/KSA/13 | KJ650297 | 160 | 240 | 80 | 120 | 120 | 80 |
| Camel/Burkina Faso/CIRAD-HKU785/2015 | C/BF/15 | MG923471 | 240 | 240 | 160 | 160 | 240 | 80 |

### ChAdOx1 MERS protects mice against different strains of MERS-CoV

To investigate whether vaccination with ChAdOx1 MERS vaccination provides protection against a variety of different MERS-CoV strains, we vaccinated *balb/c* mice transgenic for human DPP4 with ChAdOx1 MERS or ChAdOx1 GFP 28 days before challenge with 10^4^ TCID_50_ of one of six diverse MERS-CoV strains (*Figure S2*) via the intranasal route. All mice vaccinated with ChAdOx1 MERS survived challenge with MERS-CoV, independent of the challenge virus used, whereas most control mice were euthanized due to >20% weight loss or poor body condition (*Figure 6A*). Four animals per group were euthanized at 3 DPI and infectious MERS-CoV titers in lung tissue were evaluated. Whereas infectious virus could be found in lung tissue of control animals, we were unable to find infectious virus in lung tissue of ChadOx1 MERS vaccinated animals (*Figure 6B*). Thus, ChAdOx1 MERS protects against a variety of different MERS-CoV strains in hDPP4 transgenic mice.

**Fig. 6.**
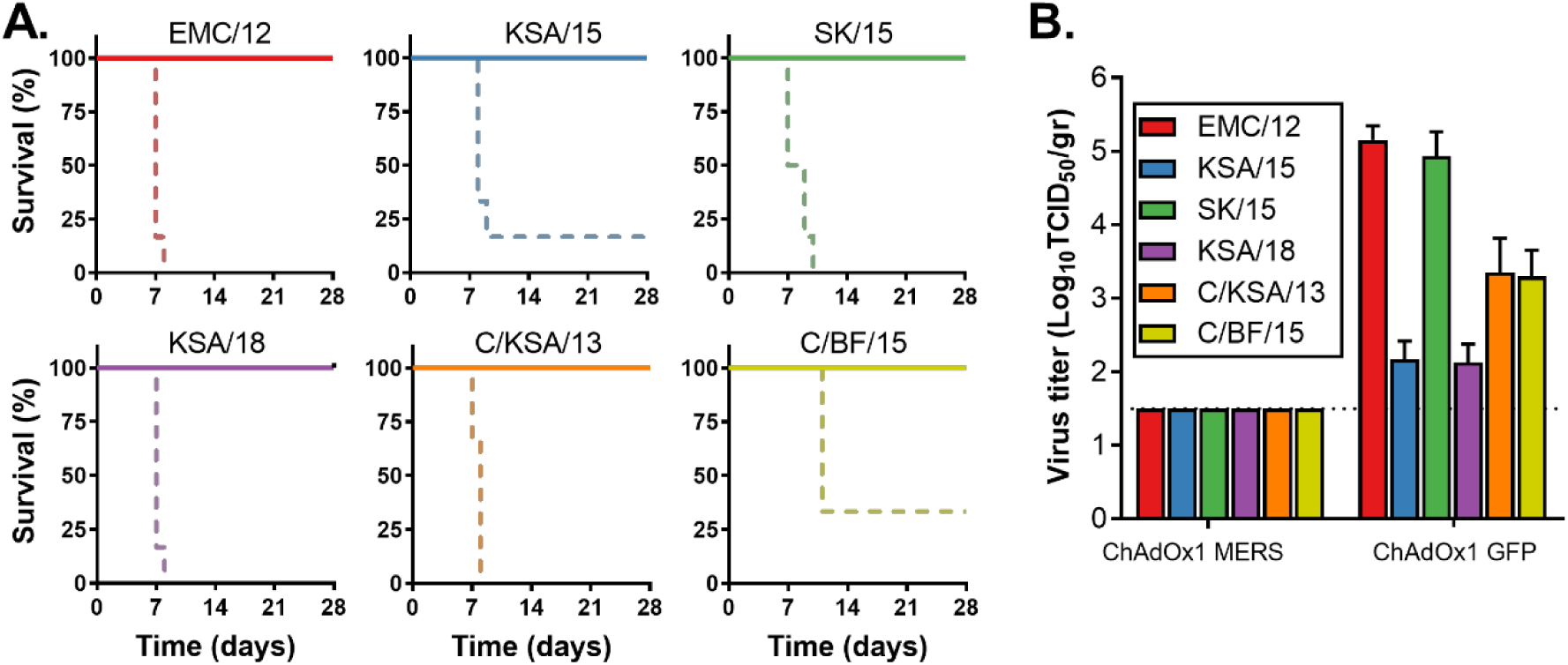
ChAdOx1 MERS provides cross-protection against different MERS-CoV strains in the mouse model. **(A)** Survival curves of ChAdOx1 MERS vaccinated (solid line) and ChAdOx1 GFP vaccinated (dashed line) hDPP4 mice challenged with MERS-CoV. **(B)** Infectious virus titers in lung tissue collected on 3 DPI from hDPP4 mice challenged with MERS-CoV. Shown is mean titer with standard deviation.

## Discussion

Middle East respiratory syndrome coronavirus is circulating in the dromedary camel population and continuously reintroduced into the human population (*17*). MERS is associated with a high case-fatality rate (34.5%) and human-to-human transmission is a major contributor to patient infections (*2*). Currently, no MERS-CoV vaccine is available. Ideally, such a vaccine would only require a single administration and would protect against a wide variety of different MERS-CoV strains.

Several studies have evaluated different types of MERS-CoV vaccines in animal models, but few have taken these vaccines into NHPs. In the current study, we show the efficacy of ChAdOx1 MERS in rhesus macaques. Unlike other NHP vaccine studies (*9-12*), we investigate vaccine efficacy after a single dose. Animals that received a single dose of ChAdOx1 MERS showed an induction of a neutralizing antibody response associated with mostly normal clinical parameters, showing no breathing irregularities or reduced lung function by spO_2_ values, limited evidence of infiltration by radiograph analysis after challenge and no signs of gross pathological lesions.

Vaccination reduced viral RNA load in tissues collected at 6 DPI compared to ChAdOx1 GFP vaccinated animals by several logs. Importantly, evidence of the presence of infectious virus in the lungs was absent and presence of mRNA in the lungs was very limited, whereas this was abundantly present in control animals. This was completely absent in animals vaccinated with a prime-boost regimen of ChAdOx1 MERS.

A variety of different MERS-CoV strains have been isolated from dromedary camels and humans over the last eight years of MERS-CoV emergence (*17*). Dromedary camels are distributed throughout Africa, the Middle East, Asia and Australia (*18*). Although MERS-CoV has not been detected in dromedary camels in Australia (*19*), strains have been isolated from Africa (*8*) and the Middle East (*6*) and seropositive dromedary camels have been found in Asia (*20, 21*). Phylogenetic analyses show a clustering of MERS-CoV by geographical location (*7, 8*) and analysis of 219 complete MERS-CoV genomes, which only included one African strain, showed the presence of two clades, with human isolates in both clades (*22*). Notably, antigenic differences have been reported between S proteins from the Middle East and Africa (*7*), potentially affecting the efficacy of a vaccine based on S protein. It is important that a MERS-CoV vaccine does not only provides homologous protection, but also against other MERS-CoV strains. We previously showed full protection of ChAdOx1 MERS (based on the Camel/Qatar/2/2014 strain) vaccinated hDPP4 transgenic mice after heterologous challenge with HCoV-EMC/2012 (*14*). Here, we repeated this challenge and extended the experiment to include five other MERS-CoV strains. We utilized strains from Saudi Arabia, South Korea and Burkina Faso, obtained from dromedary camels or humans, and isolated between 2012 and 2018 and showed full protection against all strains when mice were vaccinated with ChAdOx1 MERS in a prime-only regimen. Moreover, sera obtained at day of challenge from rhesus macaques vaccinated with ChadOx1 MERS efficiently neutralized all six MERS-CoV strains indicating that vaccination with ChAdOx1 MERS can protect against a variety of different MERS-CoV strains. Based on these results, future clinical development studies are planned with this ChAdOx1 MERS vaccine, supported by the Coalition for Epidemic Preparedness Innovations (CEPI).

In conclusion, we show that a single vaccination with ChAdOx1 MERS results in protection against disease progression and virus replication associated with MERS-CoV challenge in the rhesus macaque, and a prime-boost regimen reduced viral replication further. Furthermore, ChAdOx1 MERS vaccination protected against a diverse panel of contemporary MERS-CoV strains in hDPP4 mice. This is the first time that broad protection after a single vaccination has been shown for any MERS-CoV vaccine.

## Materials and Methods

### Ethics statement

Animal experiment approval was provided by the Institutional Animal Care and Use Committee (IACUC) at Rocky Mountain Laboratories. All animal experiments were executed in an Association for Assessment and Accreditation of Laboratory Animal Care (AALAC)-approved facility by certified staff, following the guidelines and basic principles in the NIH Guide for the Care and Use of Laboratory Animals, the Animal Welfare Act, United States Department of Agriculture and the United States Public Health Service Policy on Humane Care and Use of Laboratory Animals. Rhesus macaques were housed in individual primate cages allowing social interactions, in a climate-controlled room with a fixed light-dark cycle (12-hr/12-hr). Rhesus macaques were monitored a minimum of twice daily throughout the experiment. Commercial monkey chow, treats, and fruit were provided twice daily by trained personnel. Water was available ad libitum. Environmental enrichment consisted of a variety of human interaction, commercial toys, videos, and music. The Institutional Biosafety Committee (IBC) approved work with infectious MERS-CoV virus strains under BSL3 conditions. All sample inactivation was performed according to IBC-approved standard operating procedures for removal of specimens from high containment.

### Vaccine generation and production

The spike protein gene from MERS-CoV strain Camel/Qatar/2/2014 (GenBank accession number KJ650098.1) was codon optimized for humans and synthesized by GeneArt (Thermo Fisher Scientific). The synthesized S gene was cloned into a transgene expression plasmid comprising a modified human cytomegalovirus immediate early promoter (CMV promoter) with tetracycline operator (TetO) sites and the polyadenylation signal from bovine growth hormone (BGH). The resulting expression cassette was inserted into the E1 locus of a genomic clone of ChAdOx1 using site-specific recombination (*23*). The virus was rescued and propagated in T-REx-293 cells (Invitrogen). Purification was by CsCl gradient ultracentrifugation, and the virus was titered as previously described (*24*). Doses for vaccination were based on infectious units (IU).

### Non-human primate study

18 adult rhesus macaques (17M, 1F) were purchased from Morgan Island and randomly divided into 3 groups of six animals each. Group 1 was vaccinated with ChAdOx1 MERS at -56 DPI and -28 DPI, group 2 was vaccinated with ChAdOx1 MERS at -28 DPI, and group 3 was vaccinated with ChAdOx1 GFP at -56 DPI and -28 DPI. All vaccinations were done with 3.9 x 10^8^ IU/animal/vaccination. Blood samples were obtained before vaccination and 14 days thereafter. Animals were challenged with MERS-CoV strain HCoV-EMC/2012 on 0 DPI; with administration of 4 mL intratracheally, 1 mL intranasally, 1 mL orally and 1 mL ocularly of 10^7^ TCID_50_/mL virus solution. Clinical exams were performed on -56, -42, -28, -14, 0, 1, 3, and 5 and 6 DPI; animals were euthanized on 6 DPI. All exams existed of the following: Weight and temperature measurements, radiographs, and blood sampling. BAL was performed on 1, 3, 5, and 6 DPI by insertion of an endotracheal tube and bronchoscope into the trachea, then past the 3^rd^ bifurcation, and subsequent installation of 10 mL of sterile saline. Manual suction was applied to retrieve the BAL sample. Necropsy was performed on 6 DPI. Radiographs were evaluated and scored by a board-certified veterinarian who was blinded to the group assignment of the animals according to the following criteria: 0, normal examination; 1, mild interstitial pulmonary infiltrates; 2, moderate interstitial infiltrates, perhaps with partial cardiac border effacement and small areas of pulmonary consolidation (alveolar patterns and air bronchograms); and 3, pulmonary consolidation as the primary lung pathology, seen as a progression from grade 2 lung pathology (*25*).

### Mouse study

Groups of 10 mice were vaccinated with ChAdOx1 MERS or ChAdOx1 GFP (1×10^8^ IU/mouse) intramuscularly 28 days prior to intranasal inoculation with 10^4^ TCID_50_ MERS-CoV. Mice were challenged with one of six MERS-CoV strains: HCoV-EMC/12 (EMC/12, JX869059); Aseer/KSA-Rs924/62015 (KSA/15, KY688119); Korea/Seoul/SNU1-035/2015 (SK/15, KU308549); Riyadh/KSA-18013832/2018 (KSA/18, MN723544); Camel/Saudi Arabia/KFU-HKU1/2013 (C/KSA/13, KJ650297); or Camel/Burkina Faso/CIRAD-HKU785/2015 (C/BF/15, MG923471). Animals were monitored daily for signs of disease. Four animals were euthanized at 3 DPI and lung tissue was harvest. The remaining animals were monitored for survival. Animals were euthanized upon reaching >20% of body weight loss or poor body condition.

### Cells and virus

EMC/12 was provided by Erasmus Medical Center, Rotterdam, The Netherlands; KSA/15 and KSA/18 were provided by CDC, Atlanta, USA; SK/15 was provided by Seoul National University, Seoul, South Korea; C/KSA/13 and C/BF/15 were provided by Hong Kong University, Pok Fu Lam, Hong Kong. Virus propagation was performed in VeroE6 cells in DMEM supplemented with 2% fetal bovine serum, 1 mM L-glutamine, 50 U/ml penicillin and 50 μg/ml streptomycin. VeroE6 cells were maintained in DMEM supplemented with 10% fetal calf serum, 1 mM L-glutamine, 50 U/ml penicillin and 50 μg/ml streptomycin.

### Virus titration assay

Virus titrations were performed by end-point titration in VeroE6 cells, which were inoculated with tenfold serial dilutions of virus. After 1hr incubation at 37°C and 5% CO_2_, tissue homogenate dilutions were removed, cells were washed twice with PBS and incubated in 100 μl 2% DMEM. Cytopathic effect was scored at 5 dpi and the TCID_50_ was calculated from 4 replicates by the Spearman-Karber method (*26*).

### Virus neutralization assay MERS-CoV

Sera were heat-inactivated (30 min, 56 °C) and two-fold serial dilutions were prepared in 2% DMEM. Hereafter, 100 TCID_50_ of MERS-CoV was added. After 60 min incubation at 37 °C, virus:serum mixture was added to VeroE6 cells and incubated at 37°C and 5% CO2. At 5 dpi, cytopathic effect was scored. The virus neutralization titer was expressed as the reciprocal value of the highest dilution of the serum which still inhibited virus replication.

### Virus neutralization assay ChAdOx1

Chimpanzee adenovirus ChAdOx1 specific neutralizing antibody titers were assessed using a Secreted placental Alkaline Phosphatase (SEAP) quantitation assay. Briefly, GripTite™ MSR 293 cells (Invitrogen cat no. R795-07) were cultured as per manufacturer’s instructions and were seeded at 3×10^4^ cells per well in a 96 well plate the day prior to starting the assay (24hrs +/- 2 hrs). Cells were infected with the test sera dilutions (4 fold dilution series) at 1:18, 1:72, 1:288, 1:1152, 1:4608 in phenol red free 0% FBS DMEM (Life Technologies cat no. 31053028) and the ChAdOx1-SEAP reporter virus in a 1:1 mixture (pre-incubated for 1 hour to allow any neutralization to occur) for 1 hour before replacing with phenol red free 10% FBS DMEM for a further 24 hours (+/- 2 hrs). Sample dilutions were tested in duplicate lanes. SEAP concentration was tested on 50µl media supernatants of the samples using CPSD (Tropix PhosphaLite Chemiluminescent Assay Kit, Life Technologies cat no. T1017) using a minor variant of the manufacturer’s instructions and luminescence intensity was measured using a Varioskan Flash luminometer (Thermo Fisher). Serum dilution neutralization titers were measured by linear interpolation of adjacent values (to 50% inhibition) to determine the serum dilution required to reduce SEAP concentration by 50% compared to wells with virus alone.

### RNA extraction and quantitative reverse-transcription polymerase chain reaction

Tissues (30 mg) were homogenized in RLT buffer and RNA was extracted using the RNeasy kit (Qiagen) according to the manufacturer’s instructions. RNA was extracted from BAL fluid using the QiaAmp Viral RNA kit (Qiagen) on the QIAxtractor. The UpE MERS-CoV (*27*) or mRNA (*28*) detection assay was used for the detection of MERS-CoV viral RNA. 5 μl RNA was tested with the Rotor-GeneTM probe kit (Qiagen) according to instructions of the manufacturer. Dilutions of MERS-CoV virus stock with known genome copies were run in parallel. Genome copies were determined using Droplet Digital PCR (Biorad) and the corresponding qRT-PCR.

### Enzyme-linked immunosorbent assay

A soluble, trimeric recombinant spike protein of MERS coronavirus (isolate Ca/Jeddah/D42/2014) incorporating amino acids 1–1,273 and a carboxyl-terminal trimerization domain was produced in Chinese Hamster Ovary cells (expiCHO; ThermoFisher) and purified by immunoaffinity chromatography. Maxisorp plates (Nunc) were coated overnight at room temperature (RT) with 5 µg of S protein per plate in PBS. Plates were blocked with 100 µl of casein (Thermo Fisher) for 90 minutes at RT. Serum (2x serial diluted in casein starting at 100x dilution) was incubated at RT for 2 hrs. Antibodies were detected using affinity-purified polyclonal antibody peroxidase-labeled goat-anti-monkey IgG (Seracare, 074-11-021) in casein and TMB 2-component peroxidase substrate (Seracare) and read at 450 nm. All wells were washed 3x with PBST in between steps. Treshold for positivity was set at 3 x OD value of negative control (serum obtained from non-human primates prior to start of the experiment).

### Cytokine and chemokine profiles

Samples for analysis of cytokine/chemokine levels were inactivated with γ-radiation (2 MRad) according to standard operating procedures. Concentrations of granulocyte colony-stimulating factor (G-CSF), granulocyte-macrophage colony-stimulating factor (GM-CSF), IFN-γ, IL-1β, IL-1 receptor antagonist, IL-2, IL-4, IL-5, IL-6, IL-8, IL-10, IL-12/23 (p40), IL-13, IL-15, IL-17, IL-18 monocyte chemotactic protein-1 (MCP-1) and macrophage inflammatory protein (MIP)-1α, MIP-1β, soluble CD40-ligand (sCD40L), TGF-α, TNF-α, and VEGF were measured on a Bio-Plex 200 instrument (Bio-Rad) using the Non-Human Primate Cytokine MILLIPLEX map 23-plex kit (Millipore) according to the manufacturer’s instructions.

### Histology and immunohistochemistry

Necropsies and tissue sampling were performed according to IBC-approved protocols. Lungs were perfused with 10% formalin and processed for histologic review. Harvested tissues were fixed for a minimum of seven days in 10% neutral-buffered formalin and then embedded in paraffin. Tissues were processed using a VIP-6 Tissue Tek, (Sakura Finetek, USA) tissue processor and embedded in Ultraffin paraffin polymer (Cancer Diagnostics, Durham, NC). Samples were sectioned at 5 µm, and resulting slides were stained with hematoxylin and eosin. Specific anti-CoV immunoreactivity was detected using MERS-CoV nucleocapsid protein rabbit antibody (Sino Biological Inc.) at a 1:4000. The tissues were processed for immunohistochemistry using the Discovery ULTRA automated IHC/ISH staining instrument (Ventana Medical Systems) with a Discovery Red (Ventana Medical Systems) kit. All tissue slides were evaluated by a board-certified veterinary anatomic pathologist blinded to study group allocations.

### Statistical analyses

Tukey’s multiple comparison test or a two-tailed unpaired student’s t-test was conducted to compare differences between vaccine groups and the control group. A Bonferroni correction was used to control the type I error rate for the two comparisons (group 1 vs control and group 2 vs control), and thus statistical significance was reached at p<0.025. Spearman’s rank correlation coefficient test was used to interfere correlation.

## Acknowledgments

We would like to thank Keith Chappell, University of Queensland, Australia, for the clamped S protein used in ELISA assays, the animal care takers, Anita Mora and Austin Athman for assistance with figures, Benjamin Carrasco for preparation of animal studies, and Dana Scott, Jamie Lovaglio, Amanda Griffin, and Kathleen Cordova for assistance during the animal studies.

## Funding

This work was supported by the Intramural Research Program of the National Institute of Allergy and Infectious Diseases (NIAID), National Institutes of Health (NIH) and the Department of Health and Social Care using UK Aid funding managed by the NIHR. SCG is a Jenner investigator. The views expressed in this publication are those of the author(s) and not necessarily those of the Department of Health and Social Care.

## Author contributions

Conceptualization, NvD, TL, SCG and VJM; methodology, NvD, EH, FF, KMW, TB, RJF, AO, PWH, GS, NJE, MHAC, TL; formal analysis, NvD, and GS; writing— original draft preparation, NvD; writing—review and editing, TL, SCG, VJM.

## Competing interests

SCG is named as an inventor on a patent covering use of ChAdOx1-vectored vaccines. The remaining authors declare no conflict of interest.

## Data and materials availability

All data can be found in the manuscript and supplementary files.

## List of Supplementary Materials

**Fig. S1.**
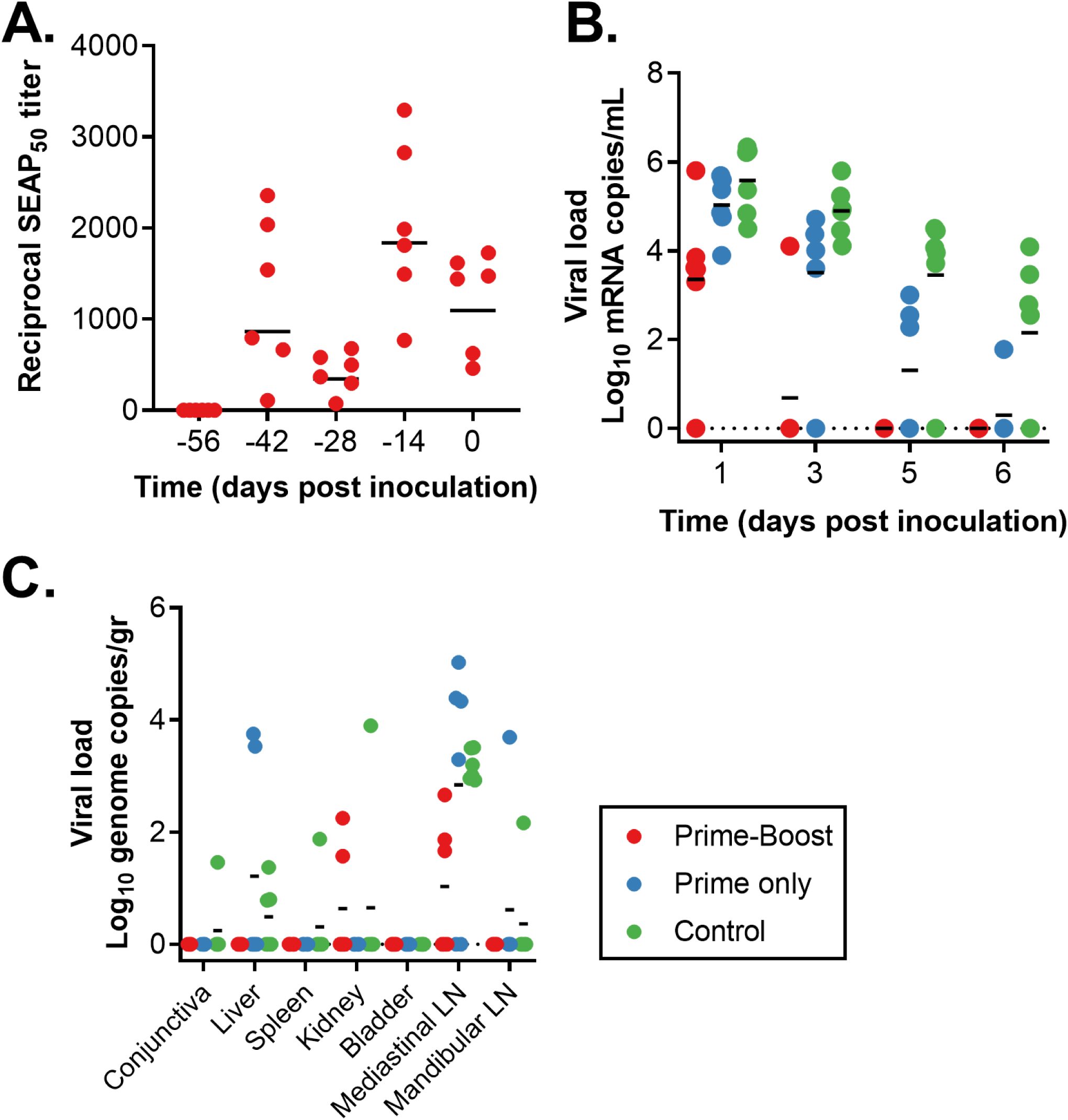
**A)** ChAdOx1-neutralizing titers in serum collected from non-human primates vaccinated via a prime-boost regimen. **B)** Viral load in BAL fluid as measured via UpE qRT-PCR. **C)** Viral load in non-respiratory tract tissue as measured via UpE qRT-PCR. Line = geometric mean. Dotted line = Limit of detection.

**Fig. S2.**
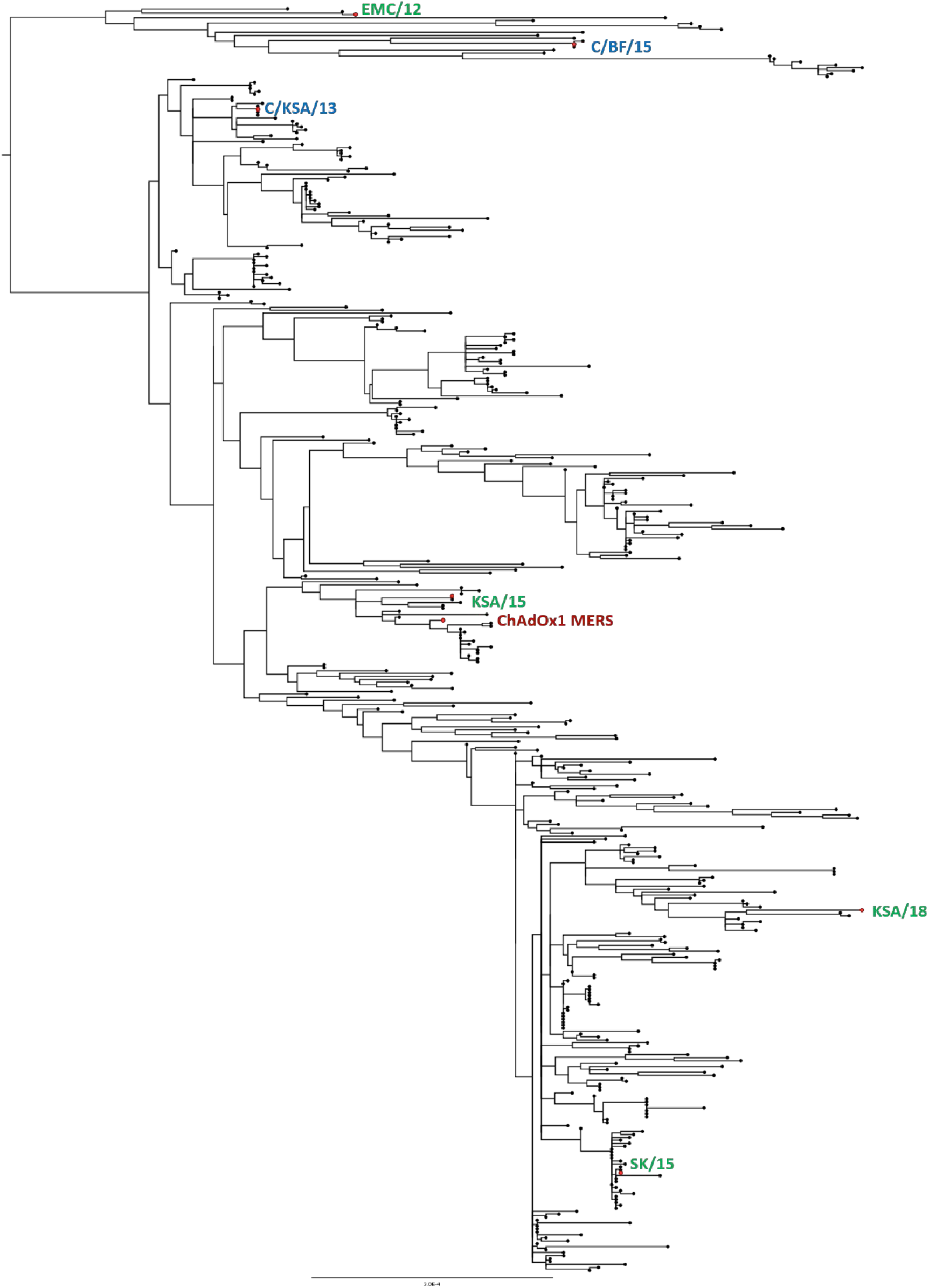
Maximum likelihood phylogeny of all published full-length MERS-CoV nucleotide sequences. Strains highlighted in green: human isolates used in this study; strains highlighted in blue: camel isolates used in this study; strain highlighted in red: ChAdOx1 MERS strain.

**Table S1.** Correlation between ELISA/VN titers and virus quantification in BAL fluid. ELISA or VN titers of all 18 animals were correlated against the amount of infectious virus, mRNA or genome copies in BAL fluid and Spearman r correlation was calculated. Spearman r correlation and two-tailed p-value shown for all combinations.

|  | Quantification method | 1 DPI | 3 DPI | 5 DPI | 6 DPI |
| --- | --- | --- | --- | --- | --- |
| ELISA | Titration | -0.69,<br>0.0015 | -0.77,<br>0.0002 | -0.53,<br>0.0230 | -0.29,<br>0.2435 |
|  | mRNA | -0.68,<br>0.0019 | -0.91,<br><0.0001 | -0.79,<br><0.0001 | -0.60,<br>0.0077 |
|  | Genome copies | -0.70,<br>0.0014 | -0.85,<br><0.0001 | -0.97,<br><0.0001 | -0.81,<br><0.0001 |
| VN | Titration | -0.69,<br>0.0015 | -0.75,<br>0.0003 | -0.53,<br>0.0248 | -0.29,<br>0.2486 |
|  | mRNA | -0.67,<br>0.0021 | -0.89,<br><0.0001 | -0.78,<br>0.0001 | -0.60,<br>0.0086 |
|  | Genome copies | -0.71,<br>0.0010 | -0.83,<br><0.0001 | -0.96,<br><0.0001 | -0.80,<br><0.0001 |

**Table S2.**
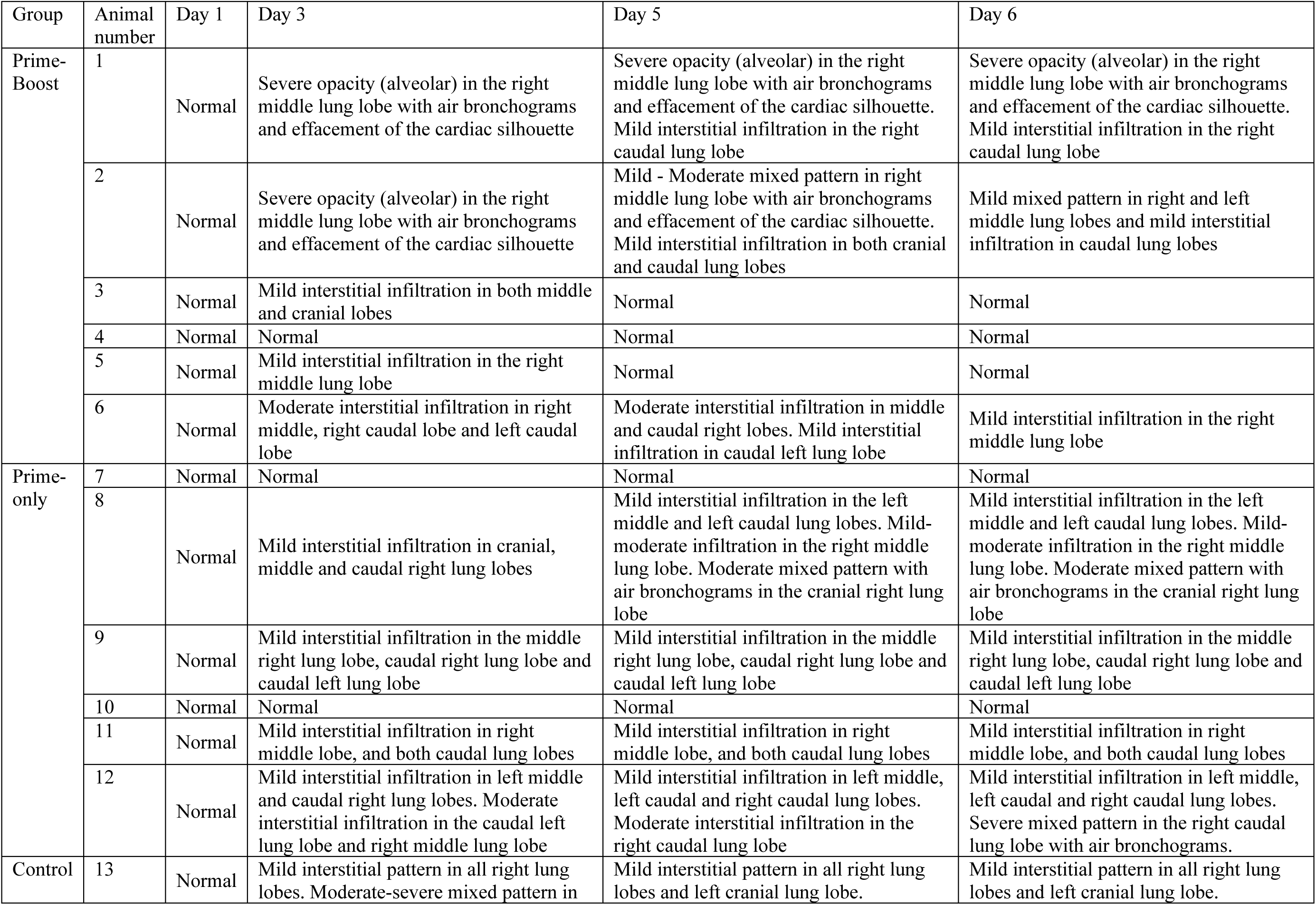
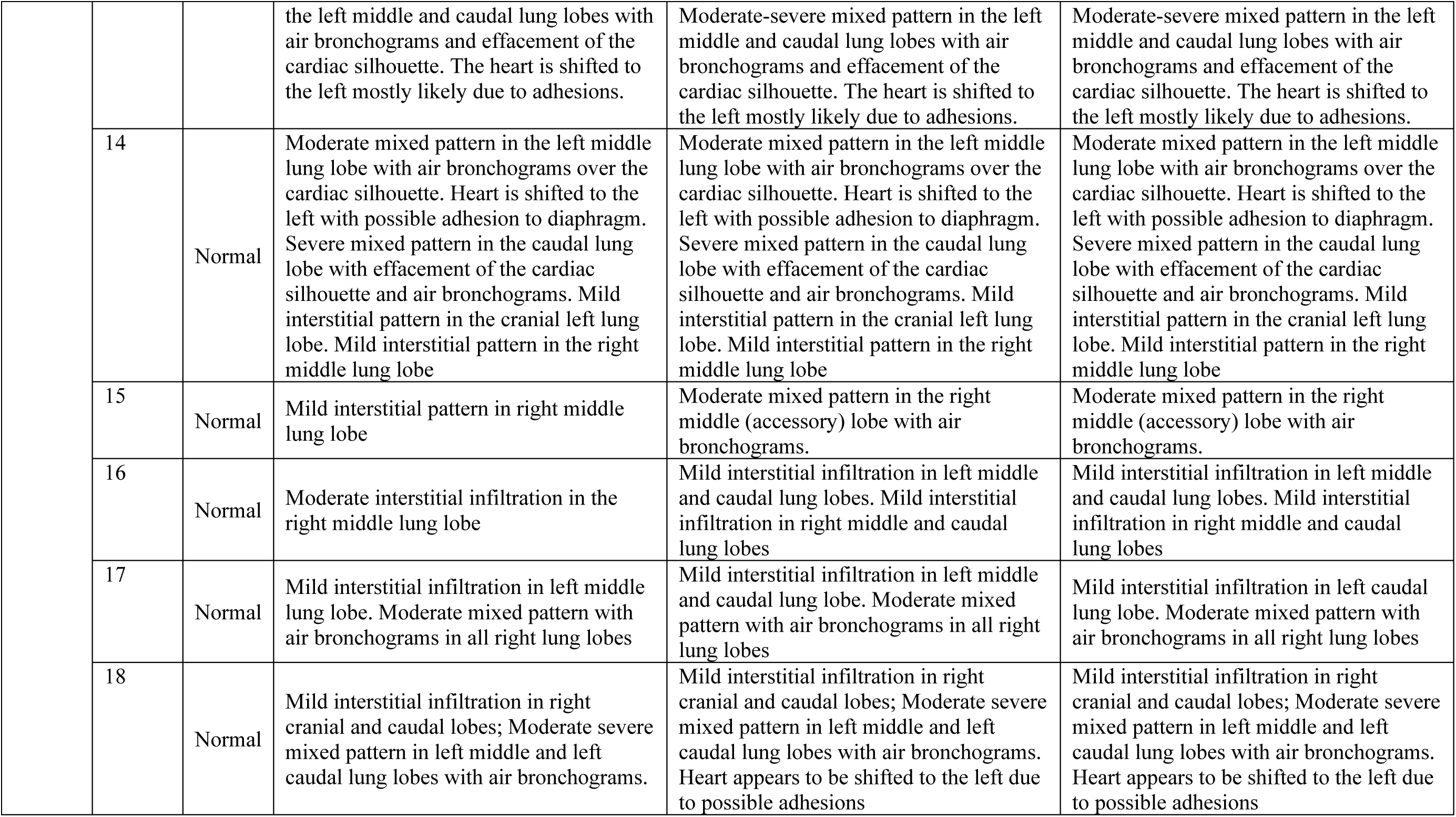
Summary of thoracic radiograph observations.

